# Fluorescent identification of axons, dendrites and soma of neuronal retinal ganglion cells with a genetic marker as a tool for facilitating the study of neurodegeneration

**DOI:** 10.1101/2024.06.20.599589

**Authors:** Puttipong Sripinun, Wennan Lu, Sergei Nikonov, Suhani Patel, Sarah Hennessy, Brent A. Bell, Claire H. Mitchell

## Abstract

This study characterizes a fluorescent *Slc17a6*-tdTomato neuronal reporter mouse line offering strong labeling in axons throughout the optic nerve, dendrites and soma in 99% of retinal ganglion cells (RGCs). The model facilitates neuronal assessment *ex vivo* with wholemounts quantified to show neurodegeneration following optic nerve crush or elevated IOP as related to glaucoma, *in vitro* with robust Ca^2+^ responses to P2X7 receptor stimulation in neuronal cultures, and *in vivo* using a confocal scanning laser ophthalmoscope (cSLO). While the tdTomato signal showed strong overlap with RGC markers, BRN3A and RBPMS, there was no cross-labeling of displaced amacrine cells in the ganglion cell layer. Controls indicated no impact of *Slc17a6*-tdTomato expression on light-dependent neuronal function, as determined with a microelectrode array (MEA), or on structure, as measured with optical coherence tomography (OCT). In summary, this novel neuronal reporter mouse model offers an effective means to increase the efficiency for real-time, specific visualization of retinal ganglion cells. It holds substantial promise for enhancing our understanding of RGC pathology in glaucoma and other diseases of the optic nerve, and could facilitate the screening of targeted therapeutic interventions for neurodegeneration.

Graphical abstract.
Fluorescent identification of axons, dendrites and soma of neuronal retinal ganglion cells with a genetic marker as a tool for facilitating the study of neurodegeneration. Puttipong Sripinun, Wennan Lu, Sergei Nikonov, Suhani Patel, Sarah Hennessy, Claire H. Mitchell*. This study delves into a new mouse model, featuring a fluorescent *Slc17a6*-tdTomato neuronal reporter. This model effectively labels axons in the optic nerve, as well as dendrites and soma in 99% of retinal ganglion cells (RGCs). This allows for both *in vitro* and *in vivo* assessment of neurodegeneration, offering a practical tool for real-time, precise visualization of RGCs, with potential applications in various fields of neuroscience and neurology. Created by Biorender.com.

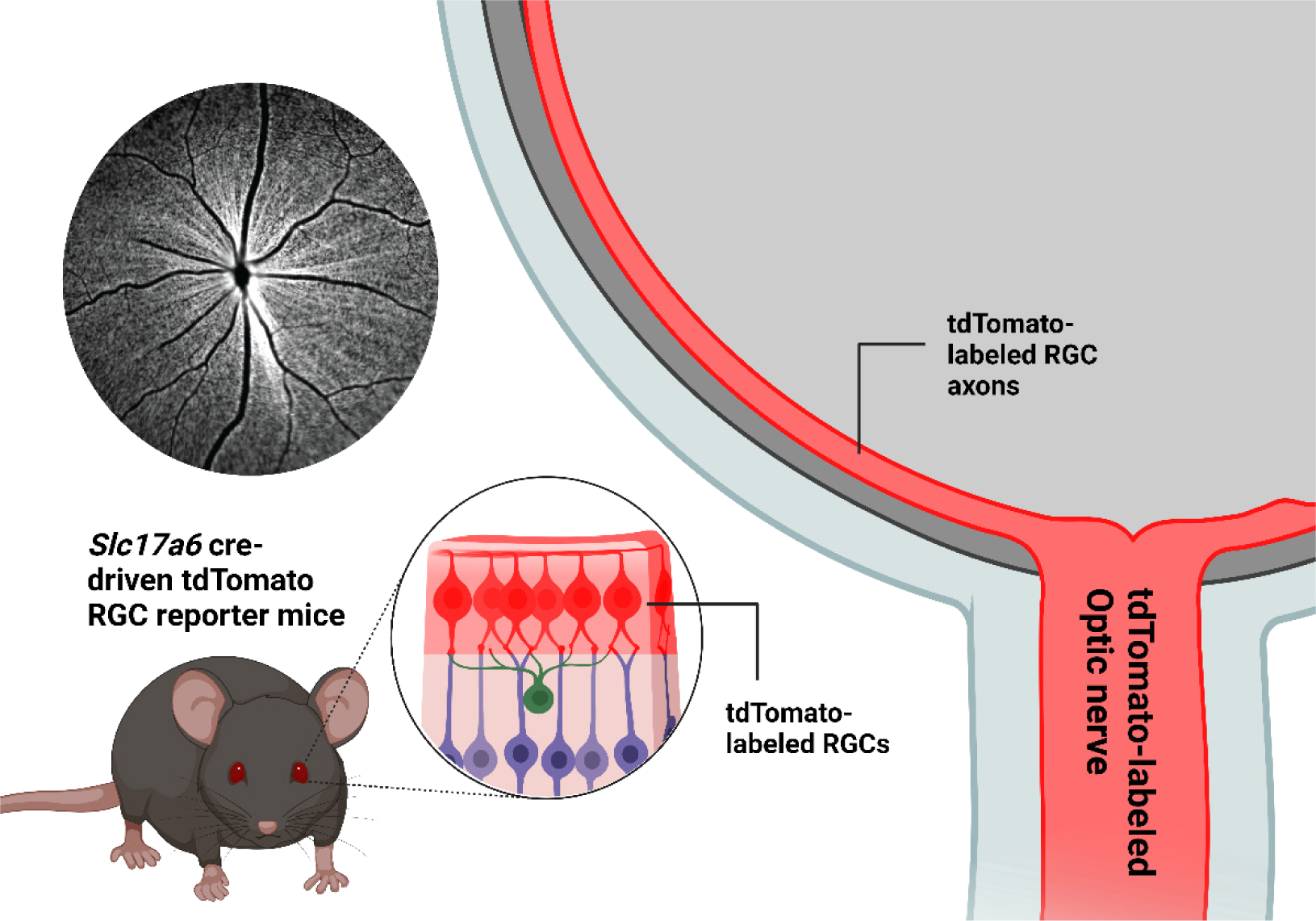

## Introduction

Retinal ganglion cells (RGCs) are an essential component of the visual system that facilitates the transmission of visual information from the eye to the brain. Functioning as the terminal neurons of the retina, they are responsible for converting the light signals detected in the outer retina into action potentials, which are then conveyed to the brain via the optic nerve ^1,2^. RGCs are particularly susceptible to damage in various ocular diseases, such as glaucoma, optic neuritis, ischemic optic neuropathy, and diabetic retinopathy ^3–6^. The rapid identification of drugs to prevent this neuronal loss requires a better way to quantify RGCs in the optic nerve and retina with advanced imaging and labeling techniques.

Several established techniques have been developed for identifying and studying RGCs, including fluorescence imaging of retrograde tracer labeling, intravitreal tracer labeling, and post-mortem immunohistochemical staining; however, each has its own drawbacks ^7^. Both retrograde and intravitreal tracer labeling, while effective for identifying RGCs, are invasive, which can cause tissue damage and inflammation due to the surgery required to deliver the tracer. These methods also can be toxic from the chemical and may result in non-specific labeling, as cell types other than RGCs, especially amacrine cells, can take up the tracer. Additionally, the tracer may only reach some RGCs, leading to incomplete labeling. These limitations highlight the need for careful consideration in selecting the appropriate labeling method to balance adequate labeling and minimize potential harm to the retinal tissue and cells.

The advent of *Thy1*-XFP reporter mice by Feng et al. ^8^ to express XFPs-fluorescent proteins, green (GFP), yellow (YFP), cyan (CFP), and red (RFP), in RGCs is beneficial for longitudinal study and can be readily detectable by non-invasive *in vivo* retina imaging. It should be noted that although *Thy1* is generally used as a specific marker for RGCs, the levels of XFP expression on the *Thy1* promoter can be quite variable in RGCs or limited to certain subtypes, depending on the integration sites. For instance, *Thy1*-YFP and *Thy1*-CFP are expressed in up to ∼ 60-80% select RGCs ^9–12^. Furthermore, the transgene has also been reported to express on some amacrine, bipolar, and Müller cells, which complicates the quantification of RGCs ^8,11,13,14^. Recently, CRISPR-Cas9 modified *Brn3b*-mCherry mice and *RBPMS*^CreERT2-tdTomato^ were developed to reduce the not specificity; however, a comparable or even smaller percentage of RGCs were labeled ^15,16^. Among others, *Slc17a6* driven-promotor has been prominently demonstrated as an excellent candidate for pan-RGCs populational studies ^17–20^. However detailed characterization studies to determine its full potential use as a RGC reporter are lacking.

In this study, we identified and systematically characterized the *Slc17a6*-driven tdTomato transgenic mice as a promising new candidate for RGC study in both *in vitro* and *in vivo* settings. We analyzed the distribution of tdTomato fluorescence within the retina and investigated its impact on RGC physiological functions. Additionally, we applied two disease models to demonstrate how this transgenic line could be used to study the course of RGC degeneration. The robust expression of tdTomato in axons, dendrites and soma of RGCs will provide a powerful tool for non-invasive observation.

## Methods

### Animal care and use

All experimental protocols were performed in accordance with the NIH Guide for the Care and Use of Experimental Animals and approved by the Institutional Animal Care and Use Committee of the University of Pennsylvania (IACUC). *Slc17a6*-driven Cre (*Slc17a6^Cre+/+^*; RRID:IMSR_JAX:028863) and Ai9 mice (*R26R^tdTomato^*; RRID:IMSR_JAX:007909), acquired from the Jackson Laboratory (Bar Harbor, ME, USA), were used to generate tdTomato-labeled RGCs mice (*Slc17a6^Cre+^*; *R26R^tdTomato+^*). Mice were socially housed and bred in cages with regular corn bedding under standard conditions with 12-hour day/night cycles and ambient humidity rooms, with food and water ad libitum.

### Immunohistochemistry

Transcardiac perfusion with PBS followed by 4% paraformaldehyde was performed on mice. Dissected eyes were post-fixed with 4% paraformaldehyde and cryoprotected in 30% sucrose overnight at 4°C, followed by embedding in optimal cutting temperature compound. Retinal wholemounts or sections were fixed with 4% paraformaldehyde for 10 minutes, permeabilized with 0.1% Triton X-100 and 20% SuperBlock buffer (ThermoFisher) in 0.1% PBS-T for 10-30 minutes at 25 °C, then blocked with 10% donkey serum and 20% SuperBlock in PBS-T for 1 hour, followed by primary antibodies 24-48 hours at 4°C with primary antibodies against BRN3A (1:50 dilution; Millipore Cat# MAB1585, RRID:AB_94166), RBPMS (1:200 dilution; GeneTex Cat# GTX118619, RRID:AB_10720427), Beta-tubulin III (1:200 dilution; BioLegend Cat# 801212, RRID:AB_2721321), and Choline Acetyltransferase (1:100 dilution; PhosphoSolutions Cat# 315-CHAT, RRID:AB_2492055). Retinal wholemounts or sections were then washed with 0.1% PBS-T and incubated with appropriate secondary antibodies conjugated to Alexa Fluor 488, 568, or 647 (1:500 dilution; Invitrogen). After incubation with DAPI, slides were mounted with SlowFade Gold antifade reagent (Thermo Fisher). Imaging was performed using a Nikon Eclipse confocal Ti2-microscope with NIS Elements Imaging software (Nikon v. 4.60).

### Two-photon imaging

Mice were euthanized by CO2 inhalation followed by cervical dislocation. Eyes were enucleated and dissected in Ames solution (Sigma-Aldrich) at room temperature and continuously oxygenated with a mixture of 95% O2 and 5% CO2. Retinas were flat-mounted with ganglion cells side up in the recording chamber perfused with the same solution maintained at 37°C using a TC-344C two-channel temperature controller (Warner Instruments, Holliston, MA, USA). All procedures were performed under dim-red light. Two-photon imaging using an Olympus FV1000 MPE Multiphoton Laser Scanning Microscope (Olympus, Center Valley, PA, USA) was employed. The wavelength for two-photon excitation was set at 1000 nm.

### Mouse models for retinal ganglion cell damage

Optic nerve crush surgery was performed based on standard approached ^21^. In brief, mice were anesthetized and maintained with 1.5% isoflurane throughout the procedure after receiving 5 mg/kg meloxicam. The temperature was maintained using a heating pad. Under a dissecting microscope, a small incision was made on the superolateral conjunctiva, and then a blunt dissection was completed with fine forceps (World Precision Instruments) to expose the optic nerve. The optic nerve was clamped with microforceps (World Precision Instruments) at a site approximately 1 to 2 mm behind the globe. After the crush injury procedure, neomycin/polymyxin B sulfates/bacitracin zinc ophthalmic ointment was applied to the surgical site, and mice were closely monitored for signs of discomfort. Mice were sacrificed after 7 days and RGC survival determined.

The transient elevation of IOP procedure is based on published approaches ^22^. In brief, anesthetized mice were administered with Proparacaine (0.5%) and Tropicamide (0.5-1%). One eye was then cannulated with a 33-gauge needle attached to polyethylene tubing (PE 50; Becton Dickinson) inserted into the anterior chamber, connected to a 20 ml syringe filled with sterile HBSS. IOP was increased to 58-60 mmHg by elevating the reservoir to the appropriate height. After 4 hrs, IOP was returned to baseline, the needle removed, and antibiotic ointment was applied to the cornea. Retinal tissues were isolated for histoimmunochemistry 10 days after the transient elevation of IOP.

### RGCs in mixed retinal cell culture

RGCs were cultured from mouse pups aged P4-6. Briefly, after euthanization, retinas were dissected and treated with papain (16.5U/mL; Cat# LK003176; Worthington) and DNase I (12,500U/mL; Cat# LK003170; Worthington) for 30 mins at 37°C. The cells were then triturated and plated on a 12 mm Nunc glass base dish (Cat# 150680; Thermo Scientific) coated with poly-D-lysine and mouse laminin I. RGC culture media ^23^ contained a 1:1 mix of base media of DMEM (Cat# 11960044; Gibco) and neurobasal (Cat# 21103049; Gibco), insulin (5 μ g/mL; Cat# I6634; Sigma), sodium pyruvate (1mM; Cat# 11360-070; Sigma), NS21 supplement (Cat# AR008; R&D systems), penicillin/streptomycin (1%, Cat # 15140-12; Gibco), SATO supplement (1X; in house-made) (Chen et al. 2008), L-glutamine (2mM, Cat# 25030-081; Gibco), triiodothyronine (T3, 40 ng/ml, Cat# T6397; Sigma), brain-derived neurotrophic factor (BDNF; 50 ng/mL; Cat# 450–02; Peprotech, Rocky Hill, NJ), ciliary neurotrophic factor (CNTF; 10 ng/mL; Cat# 450–13; Peprotech), and forskolin (4.2ng/mL; Cat# F6886; Sigma Aldrich). The cells were maintained in a serum-free RGC culture medium for 3-5 days before undergoing calcium imaging.

### Ca^2+^ measurement from RGCs

To determine the physiologic responses, levels of Ca^2+^ in RGCs were determined microscopically based on approaches described in detail previously ^22^. In brief, mixed retinal cultures were loaded with 5 µM Fura-2 AM (Thermo Fisher) with 0.01% pluronic F-127 at 37°C for 45 min. Cells were washed, mounted in a perfusion chamber, and visualized using a ×40 objective on a Nikon Diaphot microscope (Nikon, Melville, NY, USA). Ratiometric measurements were performed by alternating the excitation wavelength from 340 to 380 nm and quantifying emission ≥512 nm with a charge-coupled device camera (All Photon Technologies International, Lawrenceville, NJ, USA) Cells were perfused with isotonic solution containing 105 mM NaCl, 5 mM KCl, 6 mM 4-(2-hydroxyethyl)-1-piperazineethanesulfonic acid, 4 mM Na 4-(2-hydroxyethyl)-1-piperazineethanesulfonic acid, 5 mM NaHCO3, 60 mM mannitol, 5 mM glucose, and 1.3 mM CaCl2; the P2X7R agonist Benzoylbenzoyl-ATP (BzATP; Cat# B6396, Sigma) was added to the perfusate for the time indicated.

### In vivo imaging system

Non-invasive retinal imaging was conducted utilizing confocal scanning laser ophthalmoscopy (cSLO, Spectralis HRA, Heidelberg Engineering) and ultra-high resolution spectral-domain optical coherence tomography (SD-OCT, Envisu R2200 UHR, Bioptigen) as previously described ^24^. A 2:1 pupil dilation mixture of 2.5% phenylephrine and 1% tropicamide was applied to both eyes of the mouse, followed by anesthetization using intraperitoneal injection of a combination of 93-98 mg/kg ketamine (Dechra Veterinary Products, Overland Park, KS, USA) and 10-11 mg/kg xylazine (Akorn). Following general anesthesia, ocular protection against evaporative corneal desiccation during imaging involved using artificial tears (Refresh, Irvine, CA, USA) and ocular eye shields. Initially, both wide-field (WF-55° FOV) and ultra-wide field (UWF-102° FOV) cSLO images were captured on each eye, focusing on the retina ganglion cell layer/nerve fiber layer region using bluepeak autofluorescence (BAF-cSLO, 486nm excitation and 500-680 nm bandpass range for emission collection) images were acquired with the optic nerve centrally positioned. SD-OCT was then performed to examine the retinal structure with a 45° FOV (∼1.4 mm) with the optic nerve centrally positioned. After imaging processes were completed, mice received 1.5 mg/kg of atipamezole HCL to aid in recovery from anesthesia. To protect their eyes during recovery, Puralube Vet Ointment (Dechra Veterinary Products, Overland Park, KS, USA) was applied to the corneas.

### Microelectrode array (MEA) recordings

MEA recordings were conducted using established methods ^25^ on tdTomato-labeled RGCs and C57BL/6J mice dark-adapted for over two hours. Retinal sections were placed ganglion cell layer down in the MEA recording chamber (60MEA200/30iR-Ti-gr; Multi Channel Systems) of a 60-channel MEA system and mounted on the 1060i amplifier (Multi Channel Systems, Reutlingen, Germany), a gentle suction was applied to the sample through the perforations in the chamber bottom to enhance its contact with the electrodes, resulting in improved signal quality. The retinal tissue was perfused with oxygenated Ames solution (Sigma–Aldrich) maintained at 37°C, and stimulated with calibrated series of 455 nm light flashes. Each series included 10 2-second flashes of the same intensity delivered at 0.1 Hz frequency, different light intensities were used for different stimulation series. Data capture was acquired by an NI PCI-6071E DAQ board and custom software developed in LabView (National Instruments, Austin, TX, USA) and later analyzed using MATLAB-based custom coding (MATLAB, Natick, MA, USA).

### Data analysis

All data are displayed as mean ± standard deviation. Statistical analysis was performed using GraphPad Prism software version 10 (GraphPad, San Diego, CA, USA). Significant differences between the two related groups were assessed by t-tests with an alpha value of 0.05. Two-way ANOVA was performed when comparing more than 2 different variables. Results returning p<0.05 were considered significant.

## Results

### Characterization of tdTomato Expression in tdTomato-labeled RGCs using Slc17a6-driven Cre recombinase Transgenic Mouse Retina

Uniform and robust expressions of tdTomato were observed all across the flat-mount retina in both individual cell bodies and axons of the tdTomato-labeled RGC (*Slc17a6*^Cre+^; R26R^tdTomato+^) mice, while absence in control (*Slc17a6*^Cre-^; R26R^tdTomato+^)(Figure 1A). The pattern of tdTomato expression in transverse sections was characterized by the prominent expression of tdTomato fluorescence in the inner retina, primarily in cell bodies in the ganglion cell layer (GCL), axons in the nerve fiber layer (NFL), and dendrites extending into the inner plexiform layer (IPL). The specificity of tdTomato-positive cells in the GCL was confirmed with RGC-specific markers, with RBPMS colocalizing with the tdTomato signal in the soma and beta-tubulin III overlabbing with staining in the NFL (Figure 1B top). Colocalization with beta-tubulin III indicated strong fluorescence in the RGC axons throughout the optic nerve head (Figure 1B middle). Fluorescence was detected at the myelinated optic nerve portion as well (Figure 1B bottom). Closed inspection with two-photon microscopy demonstrated fluorescent detection of the dendrites extending into the IPL and NFL (Figure 1C). Some degree of fluorescent in a small population of horizontal ^20,26^ and photoreceptor cells ^20,27^ were evident (Supplermental Fig 3.; white and blue arrow respectively), as previously reported to be expressed in approximately 6% and 10% of that cell type’s total population, respectively.

**Figure 1.**
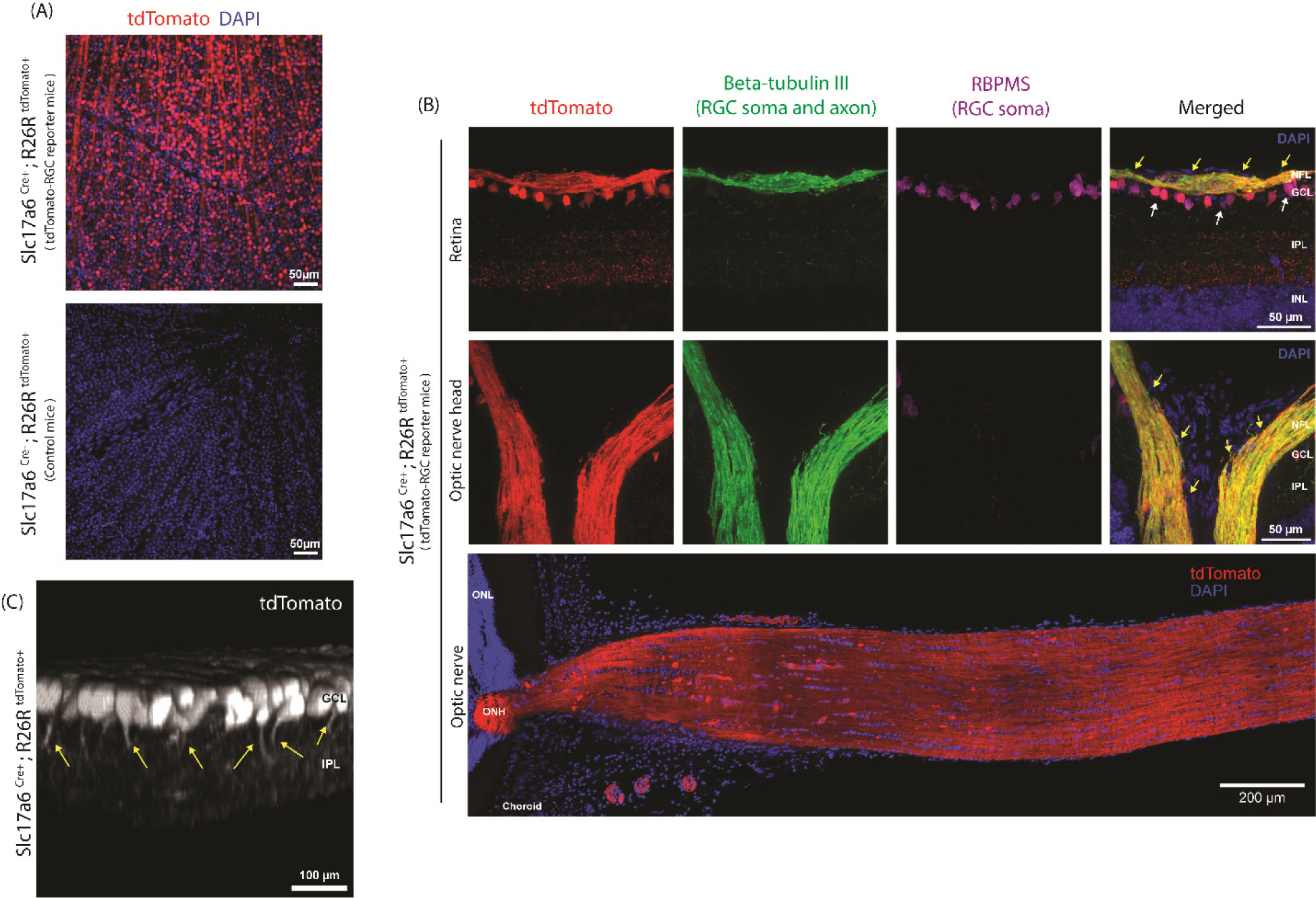
tdTomato-labeled Slc17a6-driven Cre recombinase expression characteristics in the somas, axons, and dendrites of retinal ganglion cells. (A) Cre recombinase expression drives tdTomato reporter gene in retinal ganglion cells (*Slc17a6*^Cre+^; R26R^tdTomato+^), labeling soma and axons in retinal whole mount. (B) Overlap with retinal ganglion cell markers RBPMS and beta-tubulin III confirmed tdTomato expression in retinal ganglion cells soma (top panel; white arrow) while the fluorescence overlaped with beta-tubulin III along axons continuing to the optic nerve (middle panel; yellow arrow). Fluorescent signal is detected throughout the optic nerve after it exits the globe (bottom panel). (C) A representative 2-photon imaging of tdTomato-expressing RGC promotes visualizing dendritic morphology in 3D rendering format (yellow arrow).

### Slc17a6 Cre-driven tdTomato expression labeled majority RGCs but not amacrine cells

A quantitative analysis of the relationship between tdTomato-positive cells in GCL with pan RGC-specific markers, BRN3A, and RBPMS was performed to determine the proportion of RGCs positive for tdTomato expression in the RGCs population (Figure 2A-B). All (98.93±1.4%) of RBPMS-positive cells were also labeled with tdTomato fluorochrome. Moreover, all (100±0.0%) BRN3A-positive cells also expressed tdTomato expression, which co-stained approximately 88.65±4.4% of the tdTomato-positive cells (Percentages based on analysis of 33 images from 4 *Slc17a6*-tdTomato mice stained for BRN3A and 33 separate images from the contralateral eyes of the same 4 mice stained for RBPMS). These findings indicate that the *Slc17a6* Cre-driven tdTomato is expressed by most, if not all, RGCs in consistent with previous reports which suggest that the *Slc17a6* promoter can drive Cre expression in all RGC subtypes ^18^. This contrasts with previous fluorescent mouse models only labeled a proportion of the RGCs ^8,12,14^.

**Figure 2.**
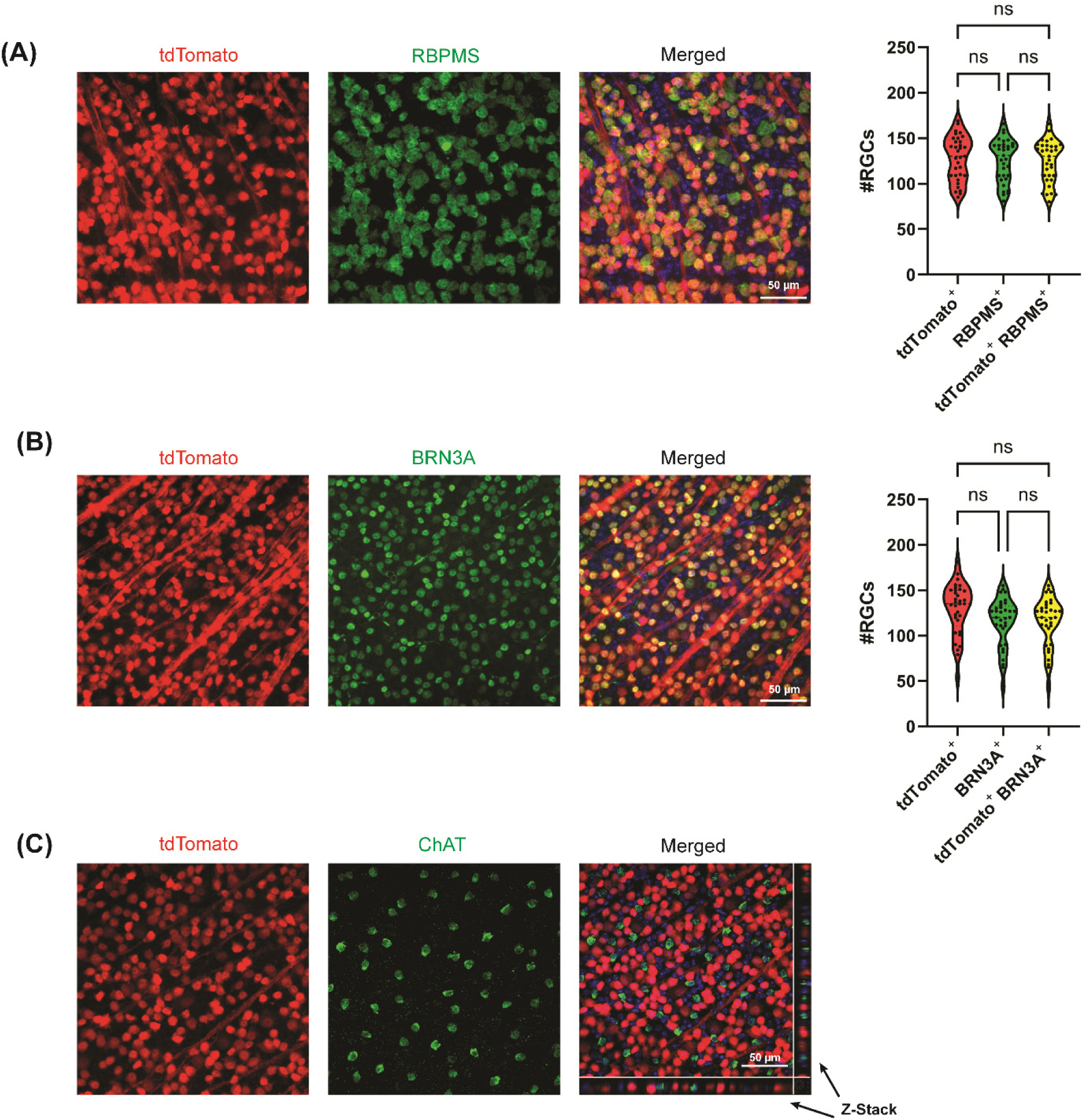
*Slc17a6* Cre-driven tdTomato expression labeled ∼99% of retinal ganglion cells but not amacrine cells. (A,B) Representative images of tdTomato (Red) and BRN3A/RBPMS (Green) showing good co-localization. Quantitative analysis of co-labeling between tdTomato-positive cells and RBPMS—or BRN3A -positive cells. (One-way ANOVA followed by Tukey post-hoc test; n = 33 images; 3 mice per group). Error bars represent mean ± SD. Statistical significance is shown as ns > 0.05, * p-value < 0.05, and ** p-value < 0.01. (C) Amacrine cells were not labeled with ChAT-positive amacrine cells showed no overlap with *Slc17a6* Cre-driven tdTomato expression in wholemount retinal sections.

Several previous reporter strains showed fluorescence in the displaced amacrine cells found among RGC bodies in GCL and complicated RGC identification ^8,11,14^. To address this, amacrine cells were identified with an antibody against choline acetyltransferase (ChAT) ^28–30^. No overlap was found between ChAT and tdTomato (Figure 2C) in agreement with previous transcriptomic ^31^ and immunofluorescence analyses ^20,32,33^, indicating that *Slc17a6* was not detectably expressed by any amacrine subtypes.

### The retina of transgenic mice expressing tdTomato under the Slc17a6 promoter can be used to follow progression in nerve injury models

Two disease models were used to inflict varying degrees of injury to RGCs. The optic nerve crush model, simulates severe injury directly to the optic nerve and is pivotal for studying the dynamics of RGC degeneration, with a significant loss of more than 60% of RGCs typically detected within a week ^34–37^. Fluorescent images from the retinal wholemount illustrate the loss of signal 7 days after the nerve crush in the affected eye as compared to the contralateral eye (Figure 3A). Rapid counting of the remaining soma indicated there was a substantial reduction of 65.92% tdTomato-labeled RGC counts seven days post-crush (Figure 3B), consistent with previous reports.

**Figure 3.**
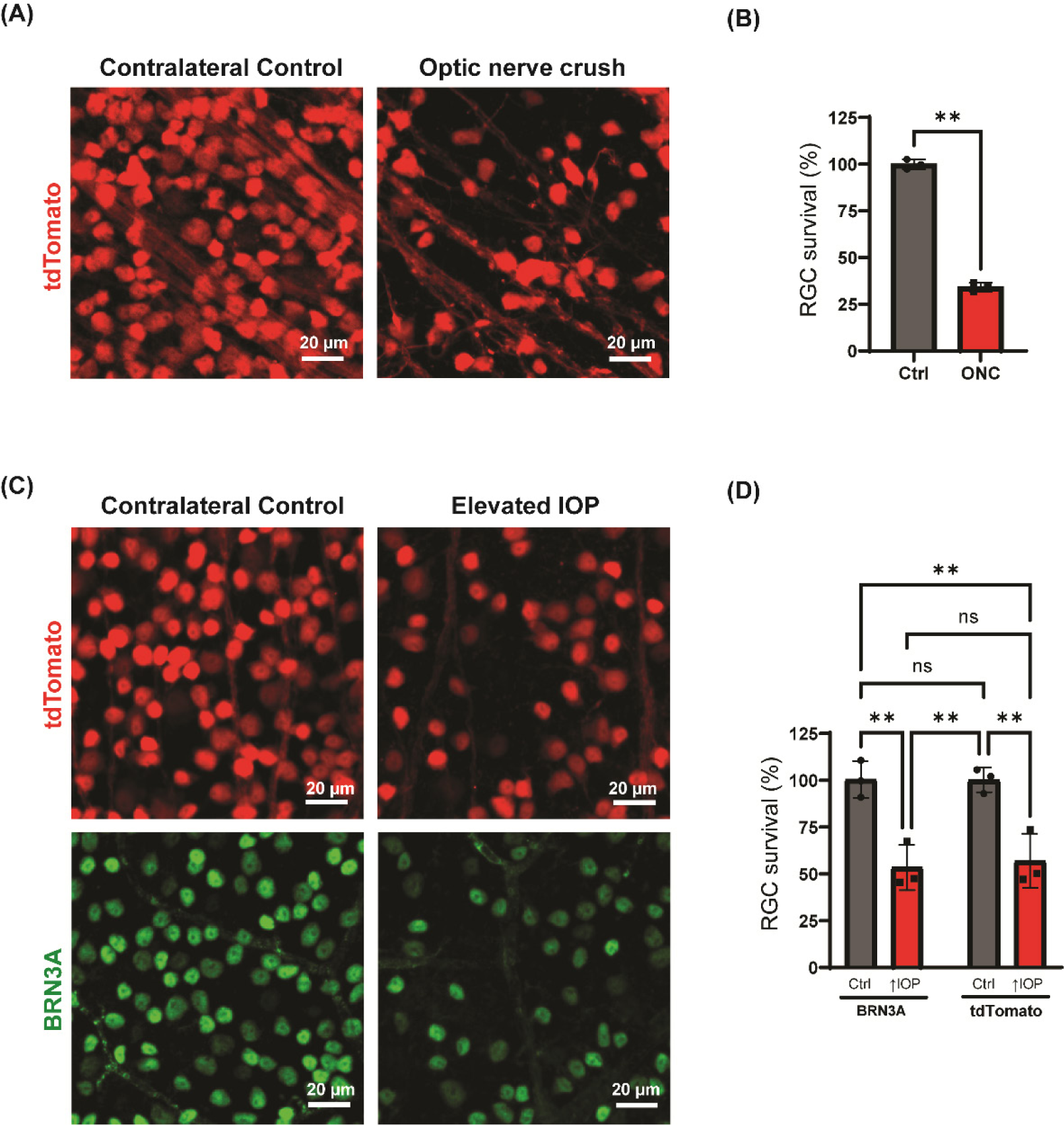
tdTomato labeling RGCs under the *Slc17a6* promoter can be used to trace the health of the RGCs. (A) Representative retinal whole mount images from the retinal wholemount of *Slc17a6*-tdTomato mice 7 days after the optic nerve of one eye one was crushed (right); the contralateral control eye is shown on the left. (B) Quantification of the RGC soma in images from the x section of mice 7 days after optic nerve crush. Significant reduction in tdTomato positive RGC soma were observed in the crushed eye compared to the control eye after 7 days. (Paired t-test; n = 3). (C) Representative retinal whole mount images from Slc7A6-tdTomato mice 10 days after transient elevation of IOP showing unlabels tdTomato mouse (top) and immunostained for BRN3A (bottom) from the same retinal wholemount. (D) Comparable reduction in BRN3A and tdTomato positive cell count in transient elevated IOP model 10 days after insult. (Two-way ANOVA followed by Tukey post-hoc test; n = 3). Error bars represent mean ± SD. Statistical significance is shown as ns > 0.05, * p-value < 0.05, and ** p-value < 0.01.

The tdTomato fluorescence was also used to track RGC survival in response to moderate transient elevation of IOP. Images indicate a loss of RGCs 10 days after the elevation of IOP, as expected (Figure 3C). Co-labelling with antibody against BRN3A of the same sections showed a similar loss. Quantifrom indicated a parallel level of RGC loss with tdTomato-labeled or BRN3A-labeled RGC counts (Figure 3D). As BRN3A is considered a gold standard marker for observing RGC health ^38^, this further corroborates the reliability of the fluorochrome detection technique in different injury contexts.

### Identification of live RGCs in primary culture of mixed retinal cells with the Slc17a6 promoter-driven tdTomato mice

Cultured neurons provide a powerful tool to examine mechanisms of neurite outgrowth and degeneration. Although neurons grow best in the presence of astrocytes, separating the cell types is complex, particularly in live cell experiments which precludes the use of antibodies. A primary culture of mixed retinal cells was developed from mouse pups to test the ability of the fluorescence to remain (Figure 4A). Strong expression of tdTomato was detected in cells from the cultures (Figure 4B). While the fluorescent signal was strongest from the soma, the signal was sufficiently bright to allow easy detection of neurites.

**Figure 4.**
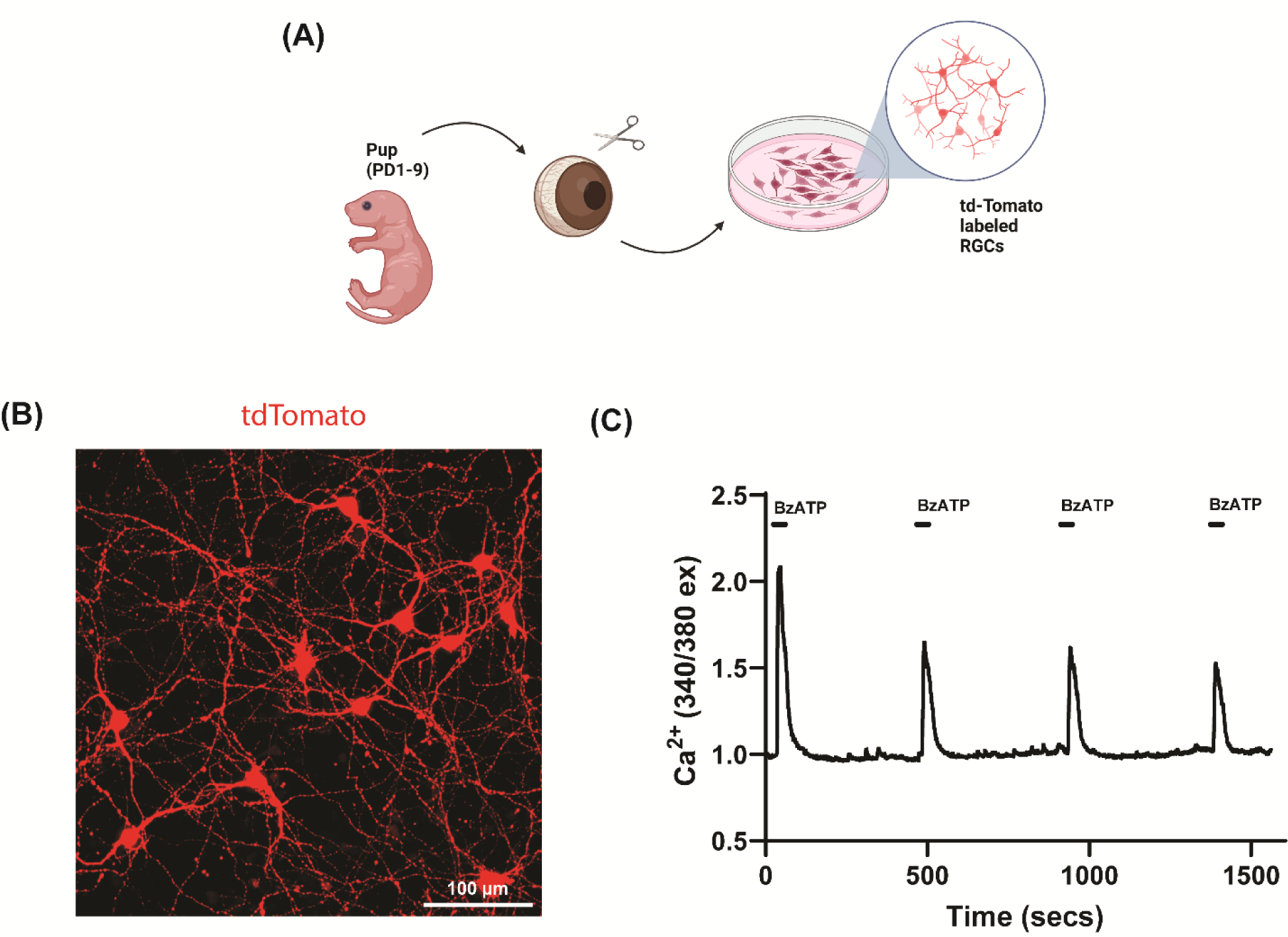
RGC identification in primary cultures of mixed retinal cells from *Slc17a6* promoter-driven tdTomato mice. (A) Schematic procedure of retinal ganglion cell isolation. (B) Representative image of tdTomato-labeled RGCs at DIV6 of mixed retinal culture. The long neurite processes can be seen extending from the soma. (C) Representative of a normalized Fura-2 Ca^2⁺^ trace of primary mouse tdTomato-labeled RGC in response to 100µM BzATP. The ratio of light excited at 340 to 380 nm provides a robust index of RGC Ca^2+^ levels from the mice.

One of the most useful assays for cultured neurons is the measurement of intracellular Ca^2+^ ; this enables receptor stimulation to be characterized and identify potential agonists to be identified. The ratiometric Ca^2+^ reporter Fura-2 was selected as its excitation at 340 and 380 nm was far removed from the tdTomato excitation of 554 nm. Red fluorescence was used to guide the selection of the “Region of Interest” from neurons. Addition of P2X7R agonist BzATP evoked rapid, reversible and repeatable responses in the RGCs identical to those recordedfrom wildtype RGCs ((Figure 4C) ^39^. This suggests that RGCs obtained from the *Slc17a6*-tdTomato are a good model to use for neuronal culture and for Ca^2+^ measurements from these neurons.

### Slc17a6 Cre-driven tdTomato allows visualization of RGCs with cSLO

The application of in vivo ocular imaging to mice is a powerful approach to advance our understanding and treatment of vision-related diseases. Non-invasive sCLO imaging was performed on the tdTomato-labeled RGCs mice to determine initial images of the baseline condition. Nerve fiber bundles could be clearly detected, with the fluorescence readout from the soma detected. (Figure 5).

**Figure 5.**
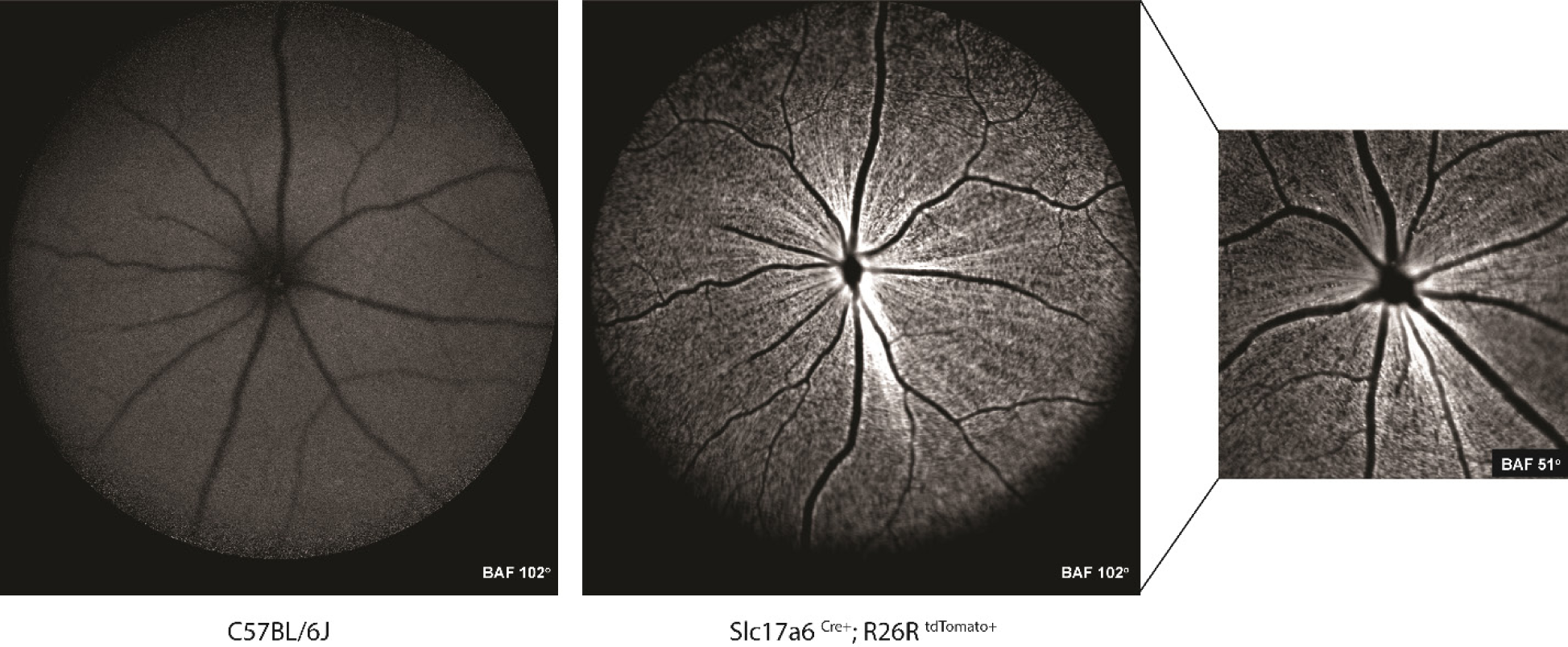
Confocal scanning laser ophthalmoscope (cSLO) allows RGC visualization in the tdTomato-labeled RGC mice in real-time. tdTomato fluorescence signals can be detected only in the retina from transgenic mice (right)but not from wild-type retina (left). Ocular diagiven in images

### Slc17a6 Cre-driven tdTomato labeling of RGCs does not interfere with RGC function or retinal structure

To confirm any gross functional changes in RGCs in the retina from the expression of tdTomato fluorochrome, Microelectrode Array (MEA) techniques were utilized. To rule out the possible effect of tdTomato on the RGC function, the electrophysiologic responses to a wide range of photopic stimuli of the retinal explants were used to compare the light-evoked firing function of the RGCs. No differences in the amplitudes of transient ON firing rate among the different groups across a range of intensities from 32 to 1.8*1e9 photons/s µm^2^, suggesting the fluorochrome does not interfere with the function of the RGCs. (Figure 6A, B)

**Figure 6.**
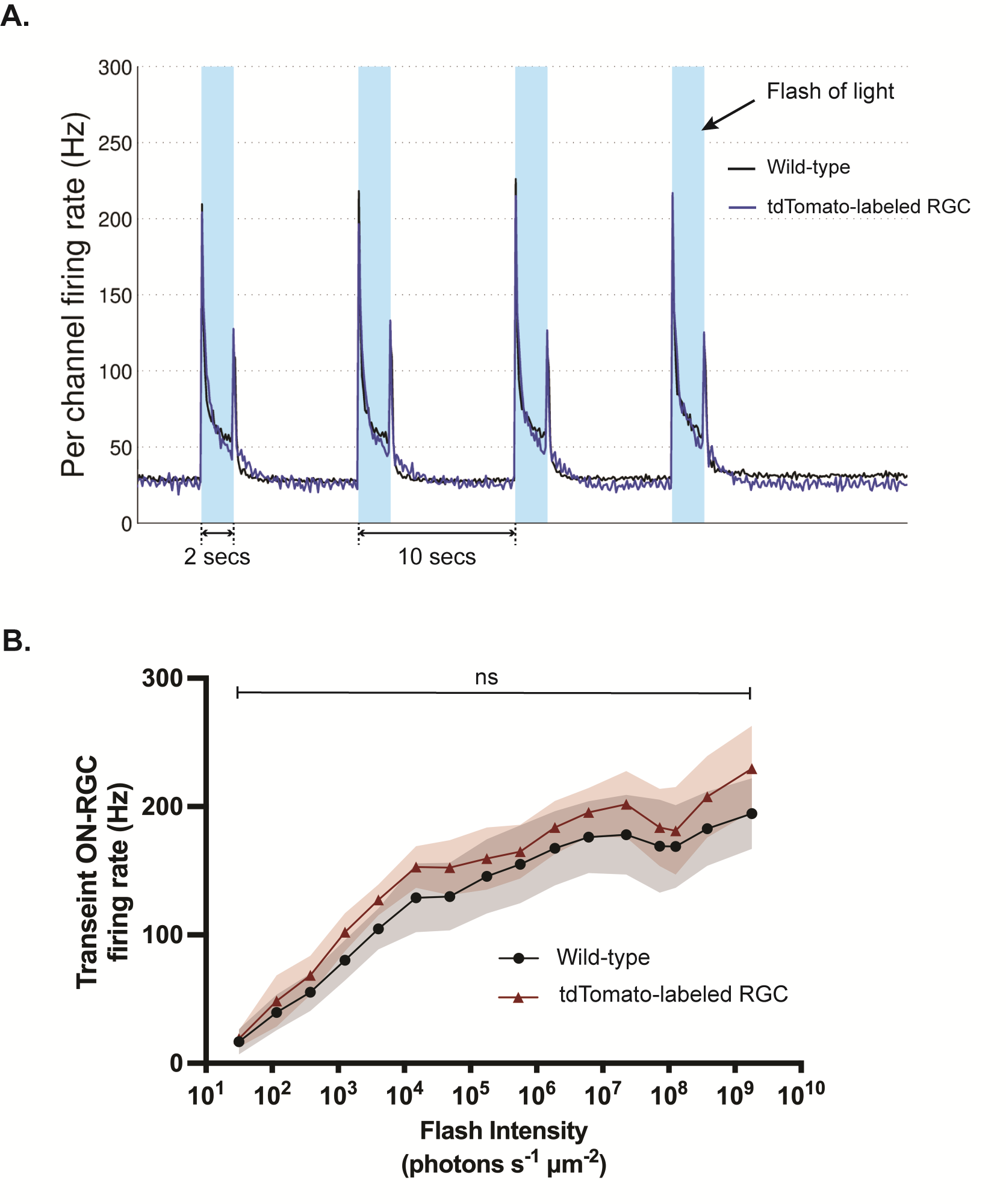
MEA analysis showed no effect of tdTomato on light-evoked RGC activity. (A) Representative MEA recording at 565,000 photons s^-1^ µm^-2^ depicted comparable light-evoked RGC firing responses between wild-type and tdTomato-labeled RGC mice; light flashes were 2 sec and 455 nm. (B) No statistical differences in the mean responses across different light intensities between wild-type and tdTomato-labeled RGC mice. (Repeated measures two-way ANOVA tests followed by Sidak post-hoc test; wild-type n = 19 retinae; tdTomato-labeled RGC n = 6 retinae). Error bars represent mean ± continuous SD. Statistical significance is shown at ns > 0.05.

To determine if expression of the tdTomato fluorochrome induced any gross pathological alteration in the retina, optical coherence tomography (OCT) was performed. OCT images were acquired at the center of the optic nerve, and retinal thickness measurements were obtained from the nerve fiber layer border to Bruch’s membrane. No statistical differences in retinal thickness were observed across various regions of the mouse retina, when retinal thickness across different distances from the optic nerve head was analyzed in a spider graph (Figure 7A, B).

**Figure 7.**
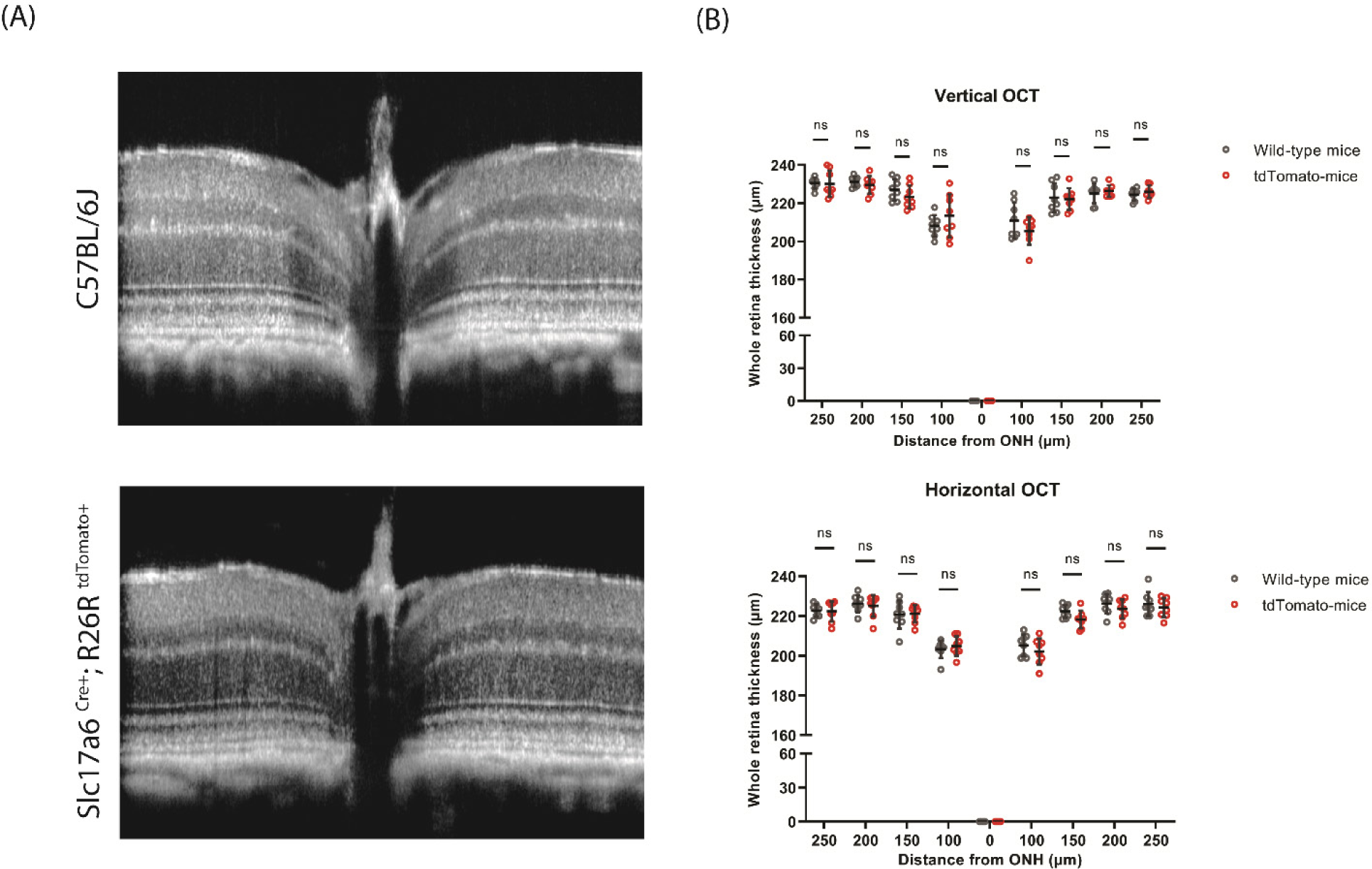
Optical coherence tomography (OCT) demonstrated normal retinal architecture and thickness in the tdTomato-labeled RGC mice. (A) Representative OCT images of C57BL/6J mouse in comparison to the tdTomato-labeled RGC mice. (B) Quantitative analyses showed a comparison of the full retinal thickness between wild-type and tdTomato-labeled RGC mice. The values in the horizontal axis of spider diagrams represent the distance from the optic nerve head where the measurements were made. (Two-way ANOVA tests followed by Tukey’s post-hoc test; n = 8 eyes; 4 mice per group). Error bars represent mean ± SD. Statistical significance is shown at ns > 0.05.

## Discussion

The present study characterizes the *Slc17a6*-tdTomato mouse line as a tool for the study of RGCs. Immunostaining with two different RGC markers confirms its precision and suggests it can label all RGCs, but not displaced amacrine cells. The intensity of the tdTomato fluorescence enabled the detection of RGC dendrites in the IPL in considerable detail when examined with 2-photon microscopy, while the expression in the axons extends beyond fluorescence detected in other models. While the red fluorescence was intense, it did not impact readout of Ca^2+^ when the UV-reporter Fura-2 was employed, nor impact RGC’s light response (light-dependent change in action potential firing frequency), as confirmed with MEA recording using 455 nm light stimuli. The *Slc17a6*-tdTomato reporter mouse offers an powerful tool to simplify the analysis of RGC survival and could increase the screening of neuroprotective agents.

### Fluorescence in dentrites, soma and axons

The intense soma labeling characteristic of the *Slc17a6*-tdTomato mouse in RGCs is comparable to that of BRN3A, which exhibits a nuclear-like pattern with well-defined borders. This similarity facilitates a more straightforward and easier quantification of tdTomato-labeled RGCs compared to other markers, such as RBPMS, which may present more diffuse or less distinct labeling patterns across the cytoplasm. Besides the soma of the RGC, a bright fluorescence is detected in several other RGC regions without the need for a primary antibody against cellular components or a fluorescence protein. The ability to detect details of dendrites projecting into the INL with the 2-photon microscope may be of help when assessing the degree of dendritic retraction associated with glaucoma ^40^. The intense fluorescence from axons in the optic nerve head and and myelinated portions of the optic nerve beyond the globe should facilitate analysis of their role in visual processing and disease, particularly as other RGC markers can show a preference for staining the soma.

The robust endogenous expression of tdTomato also significantly enhances the visualization of RGCs in an *in vivo* setting, as demonstrated by our sCLO imaging. This capability provides invaluable insights for dynamic studies of RGC physiology under pathological conditions in real-time within a living organism, with support of feasibility from the reduction in RGC numbers using immunohistochemical study in two disease models; cSLO examination with a laser more closely tuned to the tdTomato activaton spectra is expected to enhance detection further, although the intensity may require use of lower light levels than normal.

The application of tdTomato-labeled RGCs in cell culture has also demonstrated potential for monitoring the effects of molecular manipulation on neurite outgrowth and health of the RGCs, further expanding its utility in neurobiological research. Neurons grow better and are healthier in cultures that include glial cells; the presence of genetically driven fluorephore in RGCs aids in the detection of neurite growth. Mixed cultured from *Slc17a6*-tdTomato mouse are being used to detect rates of RGC degeneration *in vitro*; the ability to measure neurite length without fixation can greatly improve the temporal understanding of processes.

### Specificty and selectivity of fluorescence in Slc17a6-tdTomato mouse

The ideal reporter will fluoresce in all RGCs and only in RGCs. RBPMS expression was reported in all cells positively immunostained for melanopsin, BRN3A, and SMI-32, leading to the assumption it identified all RGCs ^41^. The analysis above showing *Slc17a6-*tdTomato mouse colocalized with 99% of the cells positive for RBPMS implies a near total *Slc17a6* of the signal to RGCs. The findings here that 87% of the fluorescent cells in the *Slc17a6-*tdTomato mouse were also positive for BRN3A is close to the proportion of RBPMS cells positive for BRN3A ^41^. This agress with the RNAseq analysis showing all types of RGCs express *Slc17a6* ^18^. The signal intensity did vary between cells; whether this reflects differences in the activity of the *Slc17a6* promotor in different RGC types is not known, although the intense signal enabled detection of fluorescence in all RBPMS positive cells. Together this implies that close to all of the RGCs are labeled in the *Slc17a6-*tdTomato mouse.

*Slc17a6* codes for vesicular glutamate transporter 2 (*Vglut2*) and *VGlut2* is extensively distributed in mouse, macaque, and human visual cortex in addition to other brain regions and somatic nerves ^42,43^. tdTomato expression is predicted to be observed in these areas as well, although this awaits confirmation for the mice used in the present study. This neural expression means crossing the *Slc17a6 Cre* mice with mice floxed for critical genes runs the risk of systemic and/or developmental issues. However, the more limited expression of *Slc17a6* in the retina makes tdTomato reporter relatively selective for RGCs in the inner retina. Although a proportion of horizontal cells were also fluorescent, spatial segregation across retail layers should provide for selective fluorescent marking.

### Advantages of the Slc17a6-tdTomato mouse over other RGC reporter transgenic mouse

RGCs represent a heterogeneous population of neurons that can be segregated into more than 40 subtypes based on their morphological, physiological, gene expression, and mosaic characteristics ^19,44,45^, thereby hindering the development of a pan-RGC reporter line. A recent comprehensive screening of 88 Cre transgenic driver lines has identified promising advancements in this area. In particular, it was found that, alongside *Thy1*-Cre, the *Slc17a6*-Cre line demonstrates significant specificity for targeting a broad range of RGC subtypes ^18^. In the present study, we demonstrated the effectiveness and specificity of the *Slc17a6*-Cre line in labeling RGC by cross-breeding with cre reporter Ai9 mice, as well as explored its potential as a valuable tool for RGC research.

Previous RGC reporter transgenic mice encountered considerable challenges due to non-specific labeling; it also inadvertently marked other cells in the GCL, such as amacrine and Müller cells ^8^. Furthermore, only 40%–45% of CFP-positive cells in GCL in *Thy1*-CFP mice were actually ganglion cells ^11^. This non-specific labeling presented a substantial drawback for studies aimed exclusively at RGCs, as it complicated the precise identification and analysis of RGCs without interference from other cell types. Our evidence suggests that *Slc17a6* Cre-driven tdTomato mouse were successfully labeled approximately 99% of the RGCs without labeling other cell types in the GCL, thus demonstrating its superior specificity and potential utility for more precise studies of RGCs.

### Expression of Slc17a6-tdTomato has no effect on gross morphology or function

While the potential benefits of the *Slc17a6*-tdTomato mouse are evident, several controls imply that the presence of a intense red fluorescence in RGCs is not itself detrimental and does not alter basic retinal health or function. The OCT analysis indicated that the thickness was not altered in the *Slc17a6*-tdTomato mouse; as retinal thickness can be altered with swelling, inflammatory response or cell loss, the lack of a change in thickness implies thet the presence of tdTomato in RGCs does not lead to any gross physiological changes. The overall retinal output, defined as the frequency of action potentials generated by the RGCs, was also the same in wildtype and *Slc17a6*-tdTomato mouse based on the MEA response. This indicates that the response to light flashes was not affected by the presence of the flurophore. Of course this could be affected by the wavelength of light used; the light flashes in our MEA system were 455 nm; whether the response to a lower frequency light is affected remains to be determined. However the results above indicate there is no impact on the overall signaling processes.

## Conclusion

The high specificity and efficacy of the reporter in real-time visualization of retinal ganglion cells underscore its potential as a powerful tool in the study of glaucoma and other neurodegenerative diseases. By providing a means to monitor RGCs *in vivo* accurately, this reporter enhances our ability to observe the progression of glaucoma and evaluate the effectiveness of potential treatments in real-time. Moreover, the successful integration of this reporter into transgenic mouse models without altering the physiological integrity of RGCs confirms its utility and reliability for continuous, longitudinal studies. Applying this RGC reporter in diverse research settings promises to broaden our understanding of RGC dynamics under various pathological conditions, leading to more targeted and effective therapeutic strategies.

## Acknowledgements

This study was supported by R01 EY015537, R01 EY015537S1, P30 EY001583.

## Author contributions

PS made substantial contributions to the conception and design of the work, the acquisition, analysis and interpretation of data, and drafted and edited the work; WL made substantial contributions to the conception and design of the work, the acquisition, analysis and interpretation of data; SN made substantial contributions to acquisition, analysis and interpretation of data; SP made substantial contributions to the analysis of data; SH made substantial contributions to the analysis of data; BAB made substantial contributions to the acquisition and analysis; CHM made substantial contributions to the conception and design of the work, analysis and interpretation of data, and drafted and edited the work. All authors have approved the submitted version and agreed both to be personally accountable for the author’s own contributions and to ensure that questions related to the accuracy or integrity of any part of the work, are appropriately investigated, resolved, and the resolution documented in the literature.

## Data Availability statement

All data generated or analyzed during this study are included in this manuscript and its supplementary information files, or are available from the corresponding author upon reasonable request.

