## Supplementary material for "Fluorescent identification of axons, dendrites and soma of neuronal retinal ganglion cells with a genetic marker as a tool for facilitating the study of neurodegeneration": Figure S1, Figure S2, Figure S3

### Supporting Information

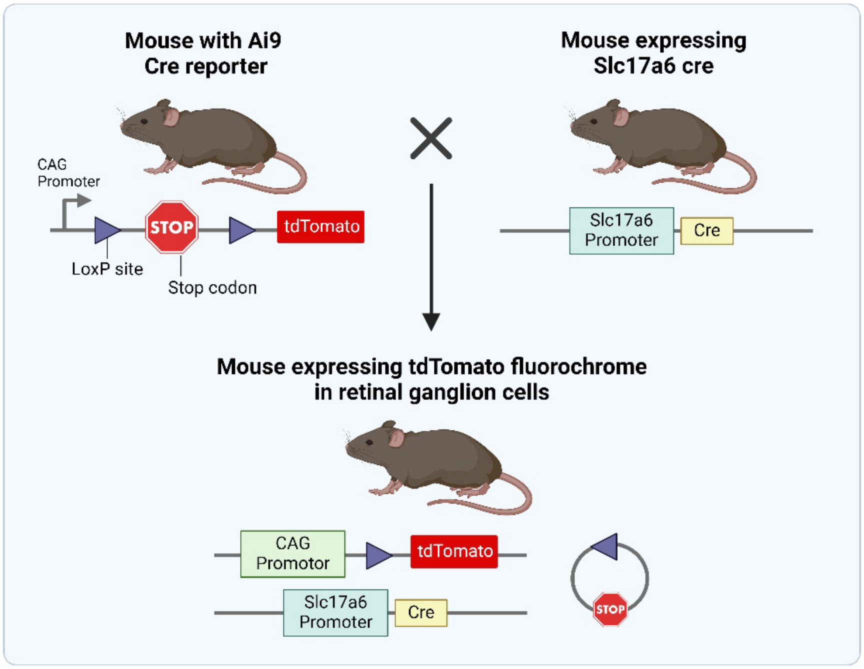


**Figure S1. Mating scheme to labeled retinal ganglion cells with tdTomato.** By crossing *Slc17a6* Cre mice with a line of mice carrying a loxP-flanked stop cassette preventing transcription of a CAG promoter-driven tdTomato, the stop cassette is excised, allowing tdTomato fluorophore expression in retinal ganglion cells. Created by Biorender.com.

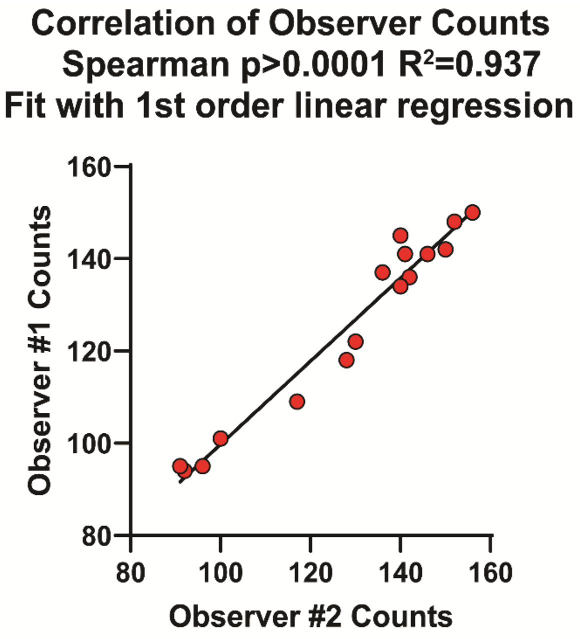


**Figure S2.** The inter-observer correlation coefficient demonstrates excellent agreement between two independent observers.

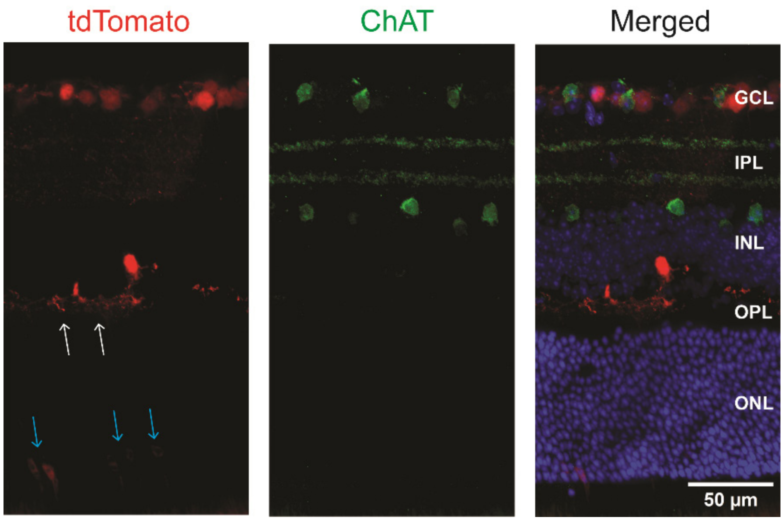


**Figure S3.** Amacrine cells were not labeled with ChAt-positive amacrine cells showed no overlap with *Slc17a6* Cre-driven tdTomato expression in transverse retinal sections. tdTomato fluorescence was also observed in a few outer retina cell types, most consistent with horizontal (white arrow) and photoreceptor cells (blue arrow).
